# Natural tolerance to transposition is associated with double-strand break repair and germ-cell differentiation

**DOI:** 10.1101/2021.04.30.441852

**Authors:** Jyoti Lama, Satyam Srivastav, Sadia Tasnim, Donald Hubbard, Savana Hadjipanteli, Erin S. Kelleher

## Abstract

Transposable elements (TE) are mobile genetic parasites whose unregulated activity in the germline causes DNA damage and sterility. While the regulation of TE mobilization by hosts is studied extensively, little is known about mechanisms that could allow germline cells to persist in the face of genotoxic stress imposed by active transposition. Such tolerance mechanisms are predicted to be beneficial when new TEs invade and host repression has not yet evolved. Here we use hybrid dysgenesis—a sterility syndrome of *Drosophila* caused by transposition of invading DNA transposons—to uncover genetic variants that confer tolerance to transposition. Using a panel of highly recombinant inbred lines of *Drosophila melanogaster*, we identified two linked quantitative trait loci (QTL), that determine tolerance in young and old females, respectively. Through transcriptomic and phenotypic comparisons, we provide evidence that young tolerant females exhibit enhanced repair of double-stranded breaks, explaining their ability to withstand high germline transposition rates. We furthermore identify the germline differentiation factor *brat* as an independent tolerance factor, whose activity may promote germline maintenance in aging dysgenic females. Together, our work reveals the diversity of potential tolerance mechanisms across development, as well as tolerant variants that may be beneficial in the context of *P*-element transposition.

## INTRODUCTION

Transposable elements (TE) are mobile DNA sequences that spread through host genomes by replicating in germline cells. Although individual TE insertions are sometimes beneficial, genomic TEs are foremost genetic parasites [reviewed in 1]. Unrestricted transposition not only produces deleterious mutations, but also double-stranded breaks (DSBs) that lead to genotoxic stress in developing gametes. Generally, hosts avoid the fitness costs of invading parasites, pathogens and herbivores by two distinct mechanisms: resistance and tolerance [2–4]. Resistance reduces parasite proliferation, whereas tolerant individuals experience reduced fitness costs from parasitism. With respect to TEs, host resistance has been the focus of extensive research, and occurs through production of regulatory small RNAs that transcriptionally and post-transcriptionally silence TEs in the germline [5–7]. By contrast, tolerance mechanisms that could ameliorate the fitness costs of transposition during gametogenesis remain largely unstudied.

The lack of research on tolerance in part reflects the ubiquity of resistance, since in the absence of high transposition rates, tolerance will not be beneficial or apparent. For example, in *Drosophila melanogaster* all actively-transposing TE families are silenced in developing gametes by the Piwi-interacting RNA (piRNA) pathway [5]. However, host genomes are frequently invaded by new TE families, against which they lack piRNA mediated resistance. Following these invasions, tolerant genetic variants may be critical for maintaining host fertility until resistance evolves. A classic example of this occured with the *P*-element DNA transposon, which invaded natural populations of *D. melanogaster* around 1950 [8–10]. When males bearing genomic *P*-elements (P-strain) are mated to naive females lacking *P*-elements and corresponding piRNAs (M-strain), they produce dysgenic offspring that do not regulate *P*-elements in germline cells [11]. A range of fertility effects result from unregulated *P*-element transposition, including the complete loss of germline cells and sterility [12]. Interestingly, naive M genotypes differ in their propensity to produce dysgenic progeny when crossed to reference *P*-strain males, suggesting the presence of tolerant variants [8,10,13,14].

In dysgenic offspring, *P*-element transposition occurs in germline cells throughout the life cycle of the fly, providing multiple opportunities for tolerant phenotypes to emerge. Starting at the second-instar larval stage dysgenic females exhibit reduced primordial germ cells (PGCs), suggesting an early onset of *P*-element transposition [15–17]. Dysgenic PGC loss is partially suppressed by overexpression of *myc*, which encodes a transcription factor that promotes stem cell maintenance [17]. PGC loss may also be suppressed by mutations in *checkpoint kinase 2 (chk2)*, a key factor in germline response to DSBs [18,19]. Tolerance of PGCs to *P*-element transposition could therefore arise through increased signaling for stem cell maintenance, or increased DNA repair in damaged PGCs.

Similar to larvae, mechanisms that reduce accumulated DNA damage, such as DNA repair, could also confer tolerance in adult females. In mature dysgenic ovaries, differentiating pre-meiotic cells undergo *chk2*-dependent cell-death at an elevated rate [15,20]. However, unlike their larval precursors the PGCs, adult germline stem cells (GSCs) are not lost at a high rate due to *P*-element transposition [15,20]. Rather, the more dramatic phenotype is that *P*-element transposition causes delay in differentiation of cytoblasts (CBs), the immediate progeny of GSCs, which results in a temporary block to oogenesis [15,20,21]. Therefore, tolerance could also emerge in adult females through mechanisms that facilitate the escape of CBs from arrested differentiation.

Through QTL mapping in a panel of highly recombinant inbred lines from the *Drosophila* Synthetic Population Resource (DSPR Population A RILs, [22]), we recently uncovered a natural tolerance allele that is associated with reduced expression of *bruno*, a female germline differentiation factor [14]. Here we present results from a second QTL mapping study in an independent panel of DSPR RILs (Population B, [22]). We describe two natural alleles that determine germline tolerance to *P*-element activity in young and aged females, respectively. We further interrogated the tolerance phenotype by contrasting RNA expression, small RNA expression, and radiation sensitivity between tolerant and sensitive genotypes, as well as by performing mutational analysis of the candidate tolerance factor *brat*. Our results suggest that young tolerant females enjoy enhanced DSB repair when compared to sensitive genotypes, allowing them to minimize dysgenic PGC loss. In contrast, we uncover the germline differentiation factor *brat* as a candidate tolerance factor in aged females. Together our results reveal the complexity of natural variation in TE tolerance, and highlight potential targets of positive selection following *P*-element invasion in natural populations of *D. melanogaster*.

## RESULTS

### 1. QTL mapping of 2^nd^ chromosome centromere

The DSPR RILs are all *P*-element free M-strains, which were isolated from natural populations before the *P*-element invasion [22]. We therefore screened for tolerant alleles among the panel B RIL genomes by crossing RIL females to males from the reference P-strain Harwich, and examining the morphology of the F1 ovaries (**Figure 1a)**. Atrophied ovaries are indicative of germline loss resulting from *P*-element activity, while non-atrophied ovaries are indicative of tolerance [14,23]. Since dysgenic females differ across development [15], and some females exhibit age-dependent recovery from *P*-element hybrid dysgenesis [24], we phenotyped F1 females at two developmental time points: 3 days and 21 days post-eclosion.

**Figure 1:**
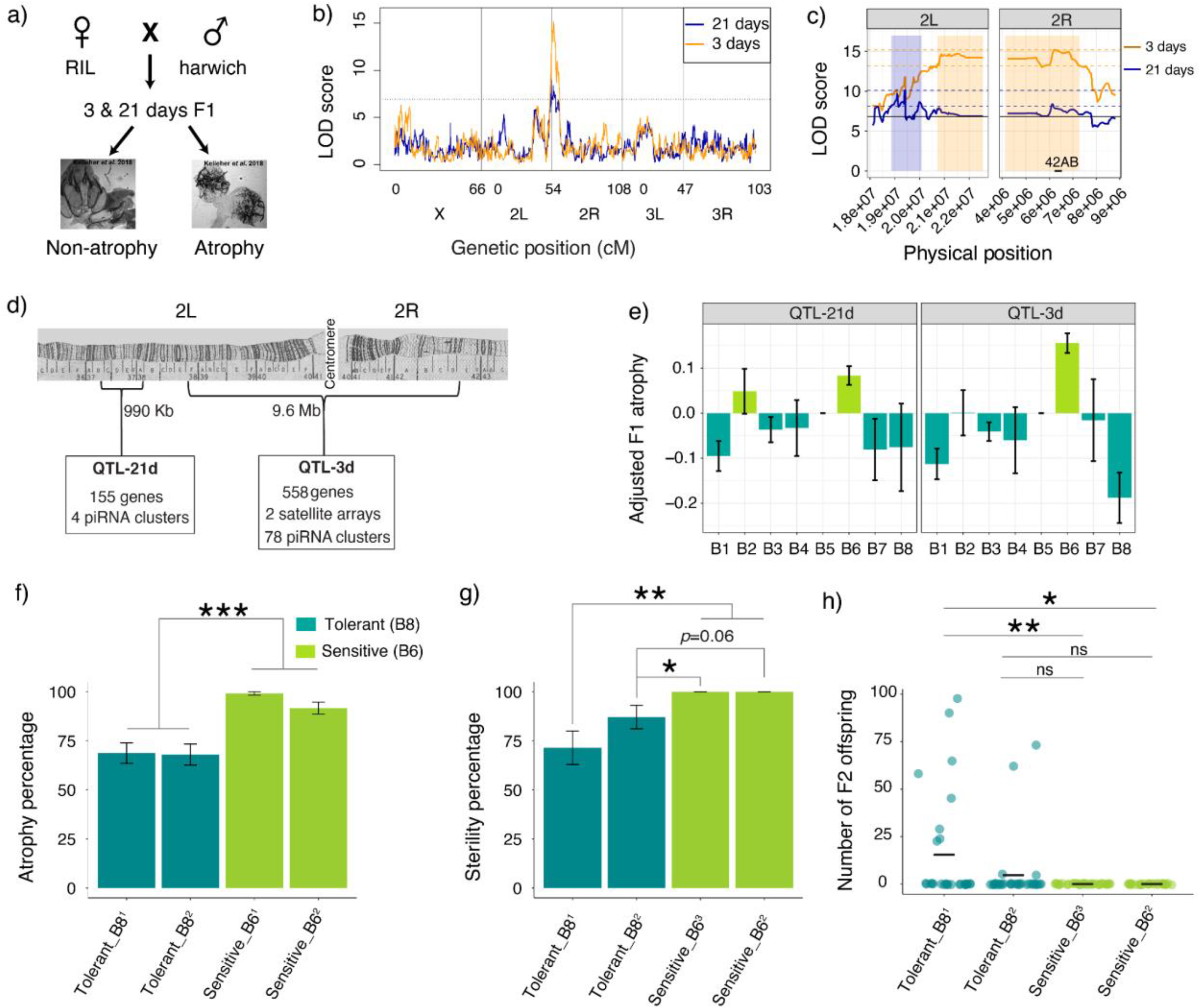
QTL mapping of variation in *P*-element tolerance. **a)** Crossing scheme to phenotype the variation in tolerance to *P*-elements among the RILs by screening for ovarian atrophy in 3 and 21 day-old dysgenic F1 females. Representative images of atrophied and non-atrophied ovaires are from Kelleher *et al*. *[14]* **b)** The log of odds (LOD) plot for QTL mapping of germline tolerance using 3 day-old (orange) and 21 day-old (blue) F1 females. The dotted line is the LOD threshold and x-axis represents the chromosomal positions. **c)** Zoomed-in figure of QTL mapping from 3 days (orange) and 21 days (blue). The colored boxes show the genomic interval that likely contains the causative genetic variant of each QTL, based on a Δ2LOD drop from the peak position [25]. The pairs of dotted lines indicate the peak Δ2LOD scores that determines the interval. The solid horizontal line is the LOD significance threshold based on 1,000 permutations of the phenotype data. **d)** Cytological map depicting the interval of the two QTL peaks [26,27]. **e)** Graph showing F1 atrophy (y-axis) associated with each of the eight founder alleles (x-axis) at the QTL peaks. All the QTL peaks show 2 phenotypic classes: sensitive (light green) and tolerant (dark green). **(f-g)** Percentage of **(f)** ovarian atrophy and **(g)** sterility among dysgenic female offspring from crosses between Harwich males and isogenic females carrying sensitive (B6) and tolerant (B8) alleles. Tolerant strains show significant reduction in F1 atrophy (Tolerant_B8^1^ vs. Sensitive_B6^1^: χ^2^= 37.05, df = 1, *p*-value = 1.15e-09; Tolerant_B8^1^ vs. Sensitive_B6^2^: χ^2^= 13.7, df = 1, *p*-value = 0.0002; Tolerant_B8^2^ vs. Sensitive_B6^1^: X-squared = 37.85, df = 1, *p*-value = 7.63e-10; Tolerant_B8^2^ vs. Sensitive_B6^2^: χ^2^ = 14.14, df = 1, *p*-value = 0.0001) as well as F1 sterility (Tolerant_B8^1^ vs. Sensitive_B6^3^: χ^2^= 10.55, df = 1, *p*-value = 0.001; Tolerant_B8^1^ vs. Sensitive_B6^2^: χ^2^= 8.41, df = 1, *p*-value = 0.003; Tolerant_B8^2^ vs. Sensitive_B6^3^: χ^2^ = 4.4, df = 1, *p*-value = 0.03; Tolerant_B8^2^ vs. Sensitive_B6^2^: χ^2^ = 3.4, df = 1, *p*-value = 0.06) compared to the sensitive strains. Subscripts ^1,2^ and ^3^ denote isogenic lines that were independently generated. **h)** Number of F2 offspring produced by individual dysgenic F1 females from crosses between Harwich males and isogenic tolerant and sensitive females. The horizontal line indicates the mean. Tolerant_B8^1^ strains show a significantly higher number of F2 offspring (Tolerant_B8^1^ vs. Sensitive_B6^3^: Z = 2.83, *p*-value = 0.004; Tolerant_B8^1^ vs. Sensitive_B6^2^: Z = 2.52, *p*-value = 0.012). Error bars in **e, f** and **g** represent the standard error. The data used to generate plot in panel **b**,**c**, and **e** are provided in **Supplemental table S3 and S4** and that used for plot in panel **f**, **g and h** are provided in **Supplemental table Supplemental figure S17** and **S18** respectively.

Similar to our observations with the Population A RILs [14], we found continuous variation in the frequency of ovarian atrophy among dysgenic offspring of different RIL mothers, indicating genetic variation in tolerance (**Supplemental table S1 and 2**). Based on a combined linear model of F1 atrophy among 3 and 21 day old females, we estimated the broad-sense heritability of tolerance in our experiment to be ~42.5%. However, the effect of age on the proportion of F1 atrophy was significant but minimal (***X***^2^= 7.03, df = 1, p-value = 0.008) with 3-day-old females showing only 0.7% increase in atrophy as compared to 21-day-old females. Therefore, age-dependent recovery from dysgenic sterility is not common among the genotypes we sampled.

To identify the genomic regions associated with genetic variation in germline tolerance, we performed QTL mapping using the published RIL genotypes [22]. We found a large QTL peak near the 2^nd^ chromosome centromere in both 3 and 21 day-old F1 females **(Figure 1b, Table 1; Supplemental table S3 and S4).** However, the genomic intervals within which the causative change separating sensitive and tolerant most likely resides are non-overlapping between the 3 and 21 day-old data sets **(Figure 1c, Table 1)**. The major QTL in 21 day-old females (hereafter, QTL-21d) resides in the euchromatic region and is quite small (990 kb) compared to the major QTL in 3 day-old females (hereafter QTL-3d), which spans the centromere and pericentromeric regions (9.6 Mb, **Figure 1d**). Therefore, there are likely at least two polymorphisms that influence tolerance near the 2^nd^ chromosome centromere, one of which is more important in young 3-day old females, and the other of which is more important in 21 day-old females.

**Table 1:** QTL positions for tolerance in 3 and 21-day old females. The peak position, Δ2LOD drop confidence interval (**Δ**2LOD CI), and the Bayesian Credible Interval (BCI) in dm6 [28] are provided for each analysis. The data used to identify the LOD peaks and intervals for 3 and 21-day old females can be found in **Supplemental table S3** and **S4**, respectively.

| Analysis | LOD Score | Peak Position | $\Delta 2$LOD CI | BCI | % variation |
| --- | --- | --- | --- | --- | --- |
| 3-day | 15.2 | 2R:6,192,495 | 2L:20,710,000-2R:7,272,495 | 2L:20,820,000-2R:6,942,495 | 11.13 |
| 21-day | 10.13 | 2L:19,420,000 | 2L:19,170,000-20,080,000 | 2L:19,010,000-20,000,000 | 9.78 |

We further evaluated the age-specific effect of two linked QTL through haplotype analysis. We modeled residual F1 ovarian atrophy as a function of QTL haplotype for the 3 day and 21 day peaks, thereby disentangling synergistic (e.g. sensitive 3d, sensitive 21d) from opposing (e.g. sensitive 3d, tolerant 21d) allelic combinations (**Supplemental figure S4**). We observed that the 3 day old QTL is solely-determinant of tolerance in the 3 day old offspring. However, in 21-day-old offspring only the genotypes containing tolerant alleles at both QTL differ from sensitive. This suggests QTL-3d may determine germ cell maintenance in the larval, pupal and early adult stages, but QTL-21d may be additionally required to maintain tolerance in aging females. The presence of two tolerance QTL is further supported by the phenotypic classes we detected among founder alleles (B1-B8) for each of the QTL peaks (**Figure 1e**). For QTL-21d, both B2 and B6 founder alleles are sensitive and greatly increase dysgenic ovarian atrophy, while all other founder alleles are tolerant. By contrast for QTL-3d, only the B6 founder allele is associated with increased sensitivity.

We next sought to determine whether reduced ovarian atrophy in tolerant alleles truly increases fitness by restoring fertility, or merely allows for the production of inviable gametes. To this end, we generated isogenic lines that carry either sensitive (B6) or tolerant (B8) alleles at both QTL loci in an otherwise identical genetic background (**Supplemental figure S5**). Consistent with our QTL mapping, tolerant alleles display less F1 ovarian atrophy (24-31%) than sensitive strains when crossed with Harwich males (**Figure 1f, Supplemental table S17**). Furthermore, fertility rates are higher than sensitive alleles (13-29)%, suggesting they are beneficial in dysgenic females (**Figure 1g, Supplemental table S18**). Finally, while tolerant females produce few offspring, offspring counts were significantly higher for tolerant females from one isogenic stock when compared to sensitive (**Figure 1h).**

### 2. Sensitive and tolerant alleles may differ in DSB repair and heterochromatin formation

Both the QTL regions contain large numbers of protein coding and non-coding RNA genes, piRNA clusters, and repeats, which could influence tolerance (**Figure 1d**). To better understand the differences between tolerant and sensitive genotypes, we compared their ovarian gene expression profiles by stranded total RNA-seq. To avoid the confounding effects of germline loss under dysgenic conditions, we focused on 3-5 day old RIL females, rather than their dysgenic offspring. To account for potential background effects, we examined three pairs of RILs that carried either a sensitive (B6) or tolerant (B4) QTL haplotype across the QTL region (dm6 2L:19,010,000-2R:7,272,495) in otherwise similar genetic backgrounds (shared 44-47% of founder alleles outside the QTL). Principal component analysis (PCA) of read counts reveals two independent axes that resolve sensitive and tolerant gene expression profiles, which together account for 40% and 16% of variation (**Figure 2a, Supplemental table S14**). One biological replicate of RIL 21188 (tolerant) was an outlier, which we excluded from our downstream analysis of differentially expressed genes.

**Figure 2:**
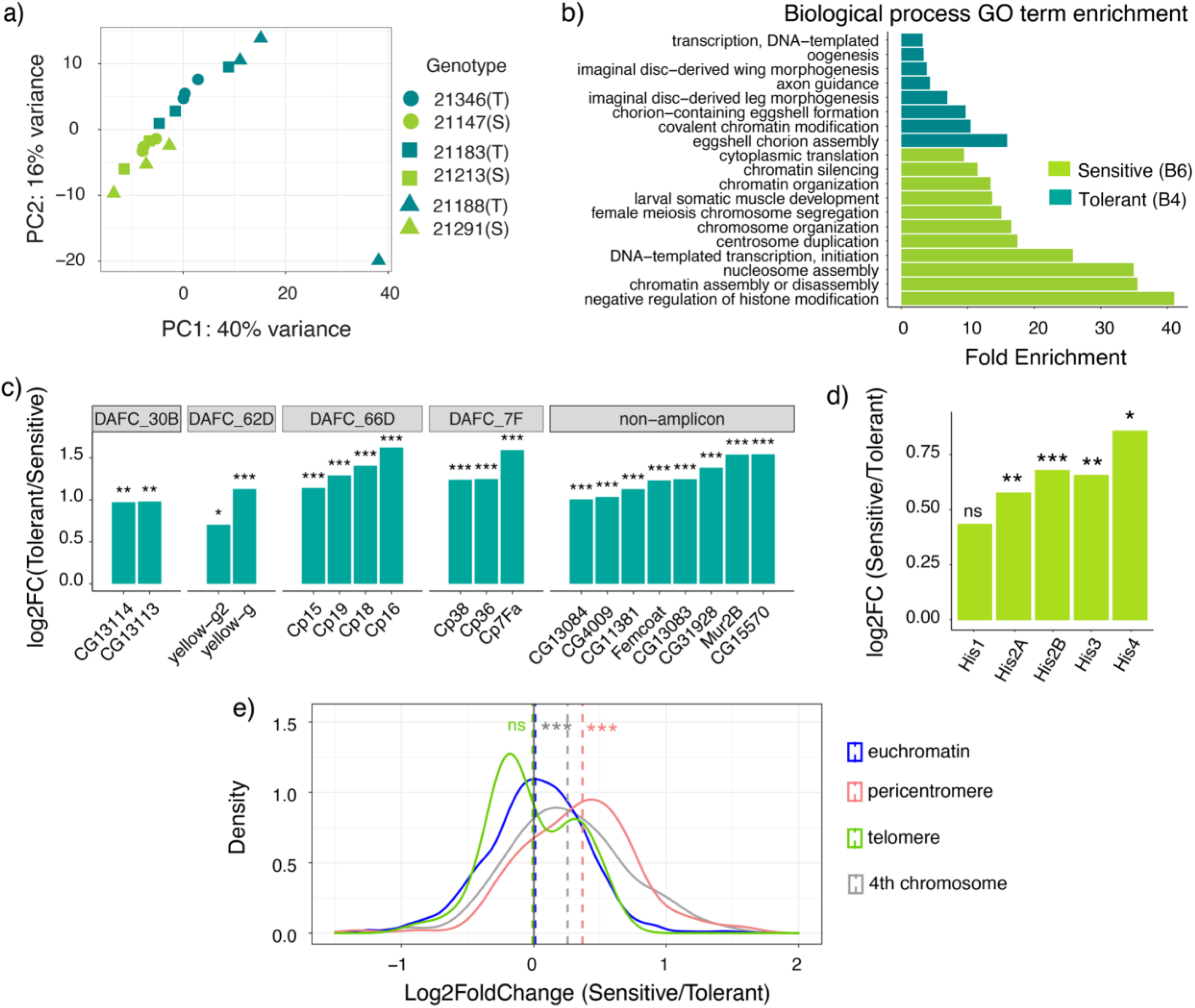
Tolerance is associated with increased chorion gene expression, whereas sensitivity is associated with increased expression of replication-dependent histones. **a**) PCA analysis of gene expression data for pairs of S/sensitive (B6) and T/ tolerant (B4) RILs. Members of the same RIL pair with otherwise similar genetic backgrounds are represented by the same shape. **b)** GO terms enriched among genes upregulated in tolerant and sensitive genotypes. **c)** Log2 fold increase in expression in tolerant genotypes for chorion genes residing in the four amplicons *(Drosophila* Amplicons in Follicle Cells, DAFCs) as well as outside amplicons [29,30]. **d)** Log2 fold increase in RD histone expression in sensitive genotypes. **e)**. Probability density plot of log2 fold change values for all euchromatic (blue), pericentromeric (red), telomeric (green) genes and 4th chromosome (gray) between strains carrying sensitive and tolerant alleles. The mean of each distribution is represented by a dotted line. Sensitive genotypes display significantly higher expression of pericentromeric genes (two-sample t-test, *t_141_*= −9.32, *p*-value = 2.335e-16) and 4th chromosome genes (two-sample t-test, *t_53_* = −4.56, *p*-value = 3.014e-05) when compared to euchromatic genes. For **e**) the x-axis boundaries were confined from (−1.5 to 2) for a better visualization. The pericentromere-euchromatin boundaries were drawn from [28,47] and subtelomeric-euchromatin boundary coordinates from [48–50]. The data represented in panel **a** is provided in **Supplemental table S14** and plot in panel **c, d,** and **e** in **Supplemental table S5**).

We found a total of 530 genes differentially expressed between sensitive and tolerant genotypes (Benjamini-Hochberg adjusted *p*-value <=0.05, fold-change > 1.5; **Supplemental table S5**). The most significantly enriched GO term among genes upregulated in tolerant ovaries is chorion assembly (Bonferroni corrected *P* value <0.01, **Figure 2b, Supplemental table S7: full report**). Indeed, all of the major chorion genes are significantly upregulated in the tolerant ovaries (**Figure 2c**, [29,30]). It is unlikely that chorion synthesis promotes tolerance because chorion synthesis occurs in late-stage oocytes [stages 10B-14, 31], whereas atrophy results from the loss of larval PGCs and pre-meiotic adult cysts (GSCs) [15–17,19]. However, chorion genes reside in clusters that undergo multiple rounds of gene amplification [32,33], generating abundant DSBs at the boundaries of the amplified region that need to be repaired to permit transcription [34]. Therefore, upregulation of chorion genes in tolerant genotypes could indicate more efficient DSB repair.

Genes upregulated in the sensitive genotypes are enriched for functions in chromatin assembly and transcription, cell division, and translation. However, a careful inspection of genes underlying these enriched terms reveals that with the exception of translation, they are majorly explained by the increased expression of replication-dependent (RD) histone gene copies (**Figure 2d**). Notably, the expression of both histone and chorion genes are increased in late oogenesis [35–38], meaning that their inverted differential expression between sensitive and tolerant genotypes cannot be explained by differential abundance of late stage oocytes. Furthermore, histone upregulation may reduce tolerance to *P*-element activity, since overexpression of RD histones is associated with increased sensitivity to DNA damage [39–43], and excess Histones are reported to compete with DNA repair proteins for binding to damage sites [40].

The *D. melanogaster* histone gene cluster is located in the pericentromeric region of QTL-3d and consists of ~100 copies of a 5-kb cluster containing each of the 5 RD histones (*his1, his2A, his2B, his3 and his4)*. However, the differential regulation of histones is unlikely to reflect the presence of a *cis*-regulatory variant within the QTL, since the histone gene cluster exhibits coordinated and dosage compensated regulation in a unique nuclear body called the histone locus body (HLB, [44]). We therefore postulate that sensitive and tolerant alleles may differ in heterochromatin formation, since many negative regulators of histone gene transcription are also suppressors of position effect variegation [43,45]. In support of this model, sensitive (B6) genotypes exhibit increased expression of pericentromeric genes, as well as genes on the heterochromatic 4th chromosome (**Figure 2e**). We also discovered increased expression of pericentromeric genes associated with the B6 haplotype in a previously published microarray dataset from head tissue ([46] **Supplemental figure S1**), suggesting B6 is unusual among the founder alleles in exhibiting reduced heterochromatin formation.

### 3. Sensitive alleles are associated with radiation sensitivity

Our gene expression data suggest that sensitive and tolerant alleles may differ in their capacity to repair DSBs. Mutations in repair genes are widely known to cause radiation sensitivity [51–55]. We therefore compared the sensitivity of the tolerant and sensitive larvae from isogenic lines to X-ray radiation.

After exploring a range of radiation doses, we found that doses above 10 Gy showed high lethality, making it difficult to detect differences in radiation sensitivity between the genotypes **(Supplemental table S19)**. Therefore, we compared the response of sensitive and tolerant larvae to radiation doses of 0 Gy, 5 Gy and 10 Gy. We observed that tolerant genotypes had significantly higher survival (53-58%) than the sensitive genotypes (25-30%) at 10 Gy (**Figure 3**). These results are consistent with differences between sensitive and tolerant alleles in DSB repair.

**Figure 3.**
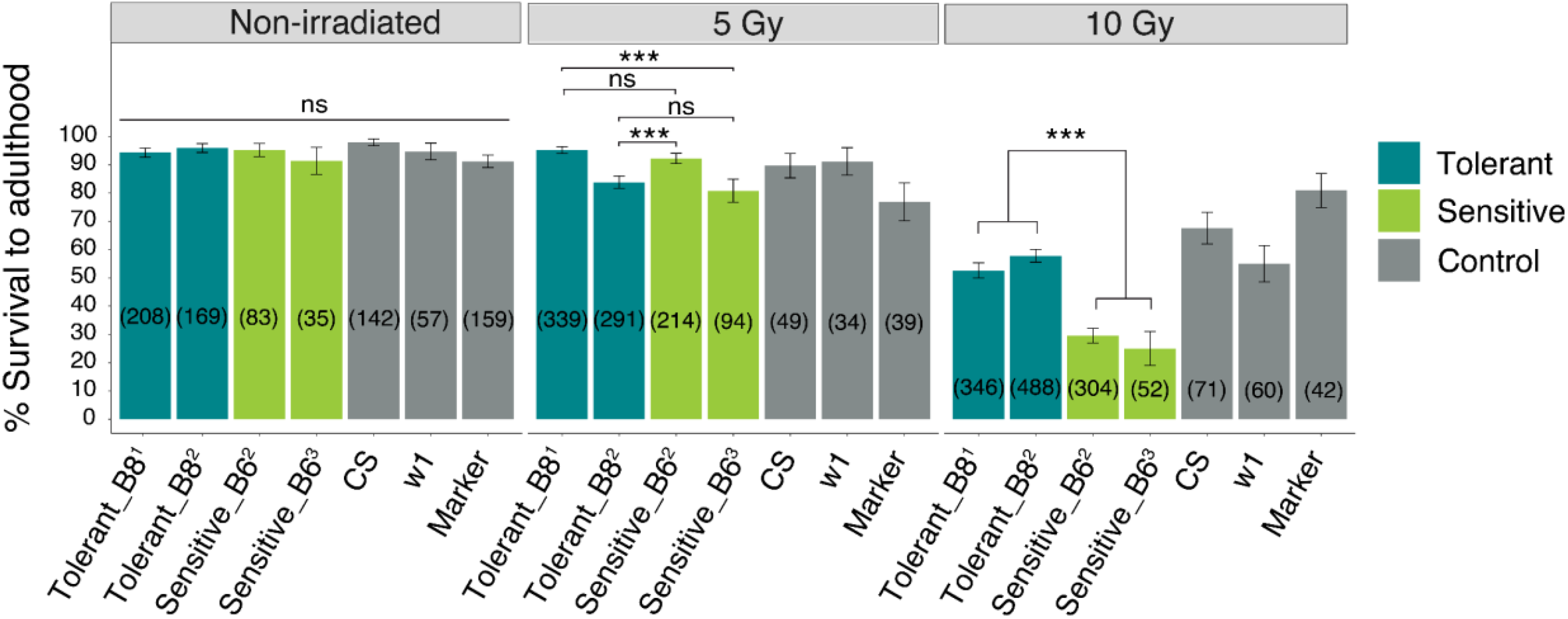
Tolerance is associated with enhanced DNA damage repair. Bar graph showing the percentage of mock treated and irradiated (5 Gy and 10 Gy) larvae that survived to adulthood for the tolerant, sensitive and the control genotypes. CS refers to Canton-S and marker refers to the multiply marked stock *b cn* (#44229), which was used to generate isogenic lines. The X-axis represents the different strains with the colors representing the type of genotype. The Y-axis is the percentage of irradiated larvae that survived to adulthood. The numbers in the brackets refer to the sample size. The number of larvae that survived and died were compared between tolerance and sensitive genotypes. For 5 Gray irradiation, Tolerant_B8^1^ vs. Sensitive_B6^3^: χ^2^= 15.66, df=1, *p*-value =0.0008; Tolerant_B8^2^ vs. Sensitive_B6^2^: χ^2^= 9.56, df=1, *p*-value =0.001. For 10 Gray irradiation, Tolerant_B8^1^ vs. Sensitive_B6^2^: χ^2^ = 34.23, df=1, *p*-value =0.0001; Tolerant_B8^1^ vs. Sensitive_B6^3^: χ^2^ = 12.69, df=1, *p*-value =0.0004; Tolerant_B8^2^ vs. Sensitive_B6^2^: χ^2^ = 58.6, df=1, *p*-value =0.0001; Tolerant_B8^2^ vs. Sensitive_B6^3^: χ^2^= 19.08, df=1, *p*-value =0.0001). The data represented in the figure is provided in **Supplemental table S19**.

### 3. piRNA clusters in QTL-3d exhibit differential activity that does not translate to TE deregulation

Although the RIL mothers do not produce or transmit *P*-element-derived piRNAs (**Supplementary table S8**), the *D. melanogaster* genome harbors >100 resident TE families [56,57] that are also regulated by piRNAs [5]. Transposition of resident TEs could add to genotoxic stress triggered by *P*-element activity, thereby reducing tolerance. Furthermore, transposition rates of resident (non *P*-element) TEs differ between wild-type strains [58–60]. Two features of our data suggest potential differences in piRNA cluster activity between sensitive and tolerant alleles. First, QTL-3d contains numerous piRNA clusters, including major ovarian piRNA cluster *42AB*, which could differ in activity between sensitive and tolerant alleles (**Figure 1d)**. Second, differential heterochromatin formation between sensitive and tolerant genotypes could impact piRNA cluster expression, which is dependent upon the heterochromatic histone modification, histone 3 lysine 9 trimethylation (H3K9me3) [61,62]. We therefore evaluated whether tolerant and sensitive alleles differ in the activity of piRNA clusters by performing small RNA-seq on the same ovarian samples used for total RNA-seq.

A PCA of piRNA cluster expression reveals that sensitive and tolerant genotypes differ in the activity of some piRNA clusters, and are resolved by the second principal component, accounting for 22% variation in expression (**Figure 4a, Supplemental table S15**). However, the major piRNA clusters—including *42AB*—are not differentially expressed between sensitive and tolerant alleles, suggesting that the proposed reduction in heterochromatin formation in sensitive genotypes does not globally inhibit piRNA biogenesis (**Figure 4b, Supplemental Table S8**). Nevertheless, we discovered two small pericentromeric piRNA clusters located within QTL-3d that were active in tolerant genotypes but largely quiescent in sensitive genotypes (**Figure 4b, c and d; Supplemental figure S2 and S3; Supplemental table S16).** These piRNA clusters are largely composed of TE fragments that are relatively divergent from the consensus (65 to 95% sequence similarity; **Supplemental table S9**), or are most similar to a consensus TE from other *(non-melanogaster) Drosophila* species. Given that transpositionally active TEs are generally highly similar to the consensus sequence [63], and piRNA silencing is disrupted by mismatches between the piRNA and its target [64], this suggests that the differential activity of these two piRNA clusters is unlikely to impact the expression of transpositionally active TEs.

**Figure 4:**
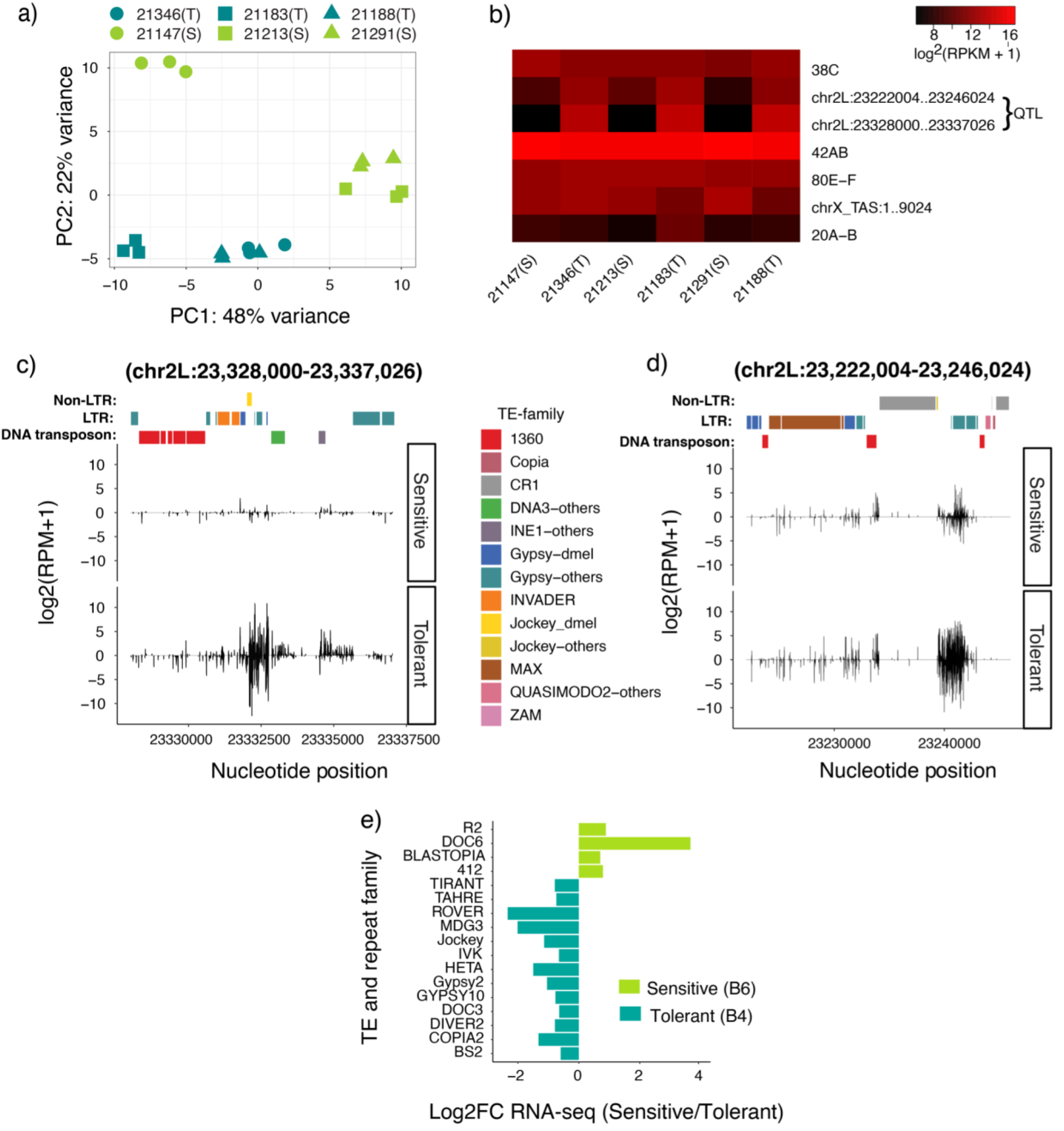
Tolerance is not determined by differential activity of piRNA cluster or TE deregulation. **a**) PCA analysis for piRNA cluster expression data of sensitive (S) and tolerant (T) genotypes. Members of the same RIL pair are represented by the same shapes. **b)** Heat map showing the expression of seven major piRNA clusters [5] and the two differentially expressed QTL clusters in QTL-3d. RIL pairs are plotted adjacent to each other. **c and d)** Uniquely mapping piRNAs within two differentially active QTL-3d piRNA clusters are compared between sensitive (21183) and tolerant (21213) genotypes. Positive value indicates piRNAs mapped to the sense strand of the reference genome and negative value indicates those from the antisense strand. TE insertions in each cluster are presented according to family by different colors; TE-others indicate the insertion was most similar to a consensus TE from a sibling species of *D. melanogaster*. See **Supplemental figure S2-3** for cluster expression in the remaining RIL pairs. For **b, c and d,** piRNA cluster expression levels are estimated by log2 scale transformed of reads per million mapped reads [log2(RPM+1)]. **e)** Genome-wide differences in TE family expression between sensitive and tolerant genotypes (fold change = 1.5, base mean >= 100, adjusted *P*-value <= 0.05), based on alignment to consensus sequences. The data used to plot panel **a** is provided in **Supplemental table S15,** for panel **b** in **Supplemental table S8**, for panel **c** and **d** in **Supplemental table S16 and S9**, and for panel **e** in **Supplemental table S10**)

To directly address if differences in tolerance are related to resident TE regulation, we compared genome-wide resident TE expression between sensitive and tolerant genotypes in our RNA-seq data. None of the TE families represented in the QTL-3d piRNA clusters were upregulated in sensitive genotypes (**Figure 4e, Supplemental table S10**). Furthermore, while some TE families are differentially expressed, there is no systematic increase in TE activity in the sensitive genotypes. Rather, more TE families are upregulated in tolerant genotypes (13 TEs) when compared to sensitive (4 TEs) genotypes. Therefore, despite the conspicuous position of QTL-3d surrounding piRNA producing-regions, as well as evidence for differential heterochromatin formation that could impact piRNA biogenesis (**Figure 2b and e**), we find no evidence that tolerance is determined by resident TE silencing.

### 4. Identifying candidate tolerance genes

We next sought to identify candidate genes that explain the tolerance differences using three criteria: 1) location within a QTL, 2) differential expression and 3) the presence of “in-phase” single nucleotide polymorphisms (SNPs) (**Supplemental table S11, S12, and S13**). In-phase SNPs are those where the genotypic differences between the founder alleles are consistent with their tolerance phenotype class [65]r**e 5a,** [65]). Of 530 differentially expressed genes (**Figure 5b**), 43 are within the QTL region, representing an approximately five-fold enrichment in the QTL regions compared to the rest of the genome *(X-squared* = 255.54, *df* = 1, *p*-value < 2.2e-16, **Figure 5b**). Ultimately, we identified 14 and 5 differentially expressed genes that also carry in-phase SNPs within the QTL-3d and 21d, respectively (**Figure 5c and d**; **Supplemental table 12**). Furthermore, we identified 37 genes in QTL-3d and 4 genes in QTL-21d containing in-phase non-synonymous SNPs, which may affect the function of the encoded protein (Supplemental table S13). These genes represent the strongest candidates to contain tolerant variants.

**Figure 5:**
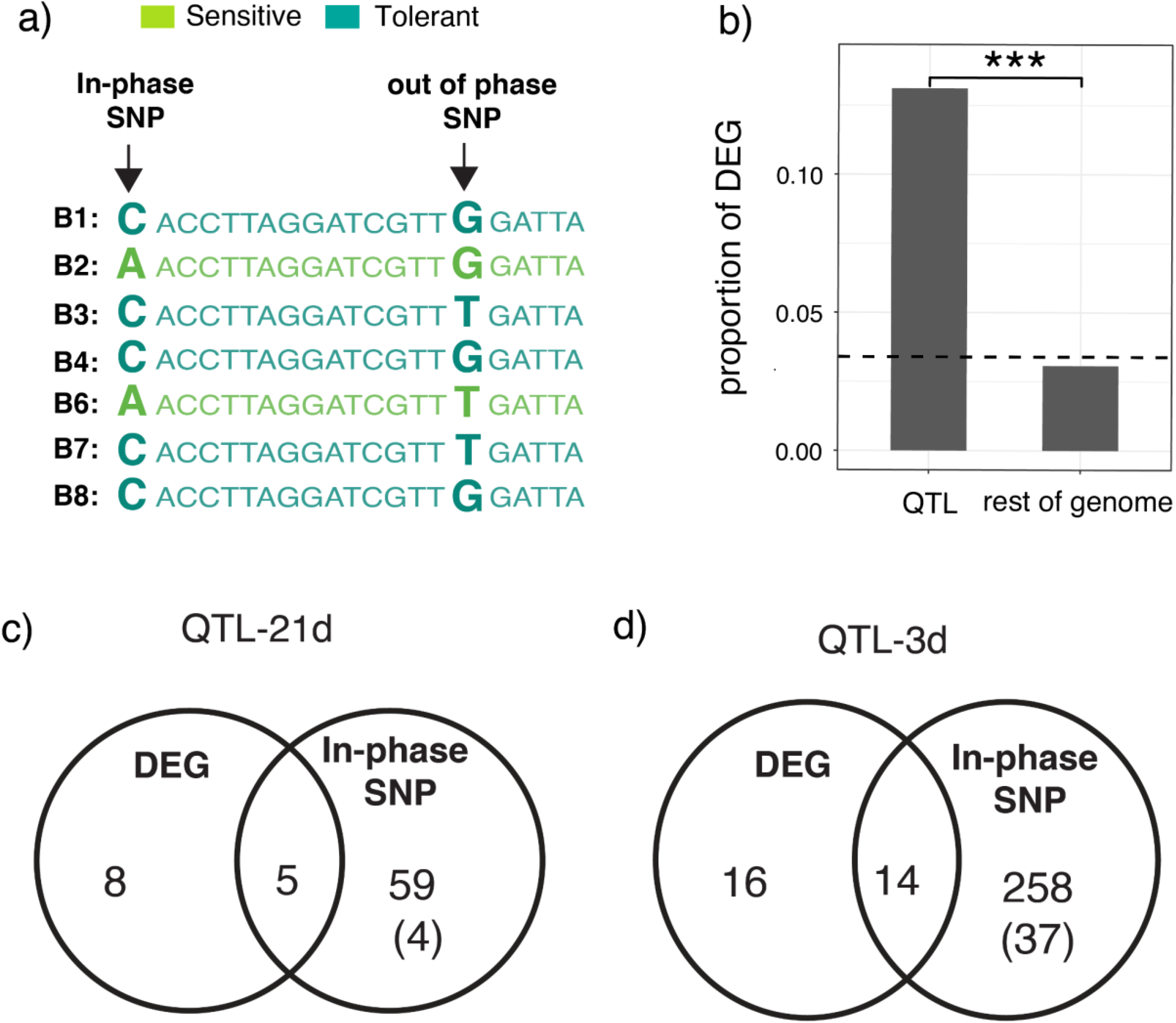
Differential expression and in-phase SNPs identify candidate tolerance genes. **a)** Hypothetical in-phase and out of phase SNPs are shown. Sequences of each of the B founder B strains are colored based on their phenotypic classification, either tolerant or sensitive (**Figure 1e**). Bold letters indicate SNPs. **b)** The proportion of genes differentially expressed (DEG) is compared inside and outside the QTL. The dotted line is the genome wide average. **c and d)** Venn diagrams showing the overlap of differentially expressed genes (DEG) and genes carrying in-phase SNPs for QTL-21d (c) and QTL-3d (d). The number outside the bracket indicates all genes with in-phase SNPs, whereas the number within the brackets indicates the genes carrying non-synonymous in-phase SNPs only. The data for differential expression of genes for tolerant and sensitive genotypes is provided in **Supplemental table S5**. The data on in-phase polymorphisms for each QTL peak are provided in **Supplemental table S11**. List of candidate genes that have both in-phase polymorphisms and are differentially expressed, and those having non-synonymous in-phase polymorphisms are provided in **Supplemental table S12** and **S13**, respectively.

We next scoured our list of candidate genes for those with known functions in heterochromatin formation and DSB repair, whose differential function or regulation are plausibly related to phenotypic differences associated with sensitive and tolerant alleles. Within QTL-3d, *Nipped-A*—which contains a non-synonymous in-phase SNP—stood out as a member of the Tat interacting protein 60 kD (TIP60) complex. The TIP60 complex has functions in DSB repair and heterochromatin formation [66–70]: providing a clear connection to our gene expression and radiation assays. The non-synonymous SNP that separates sensitive and tolerant alleles of this gene are located in the HEAT2 domain, which is predicted to be essential for protein-protein interaction [71–73]. Furthermore, two additional members/interactors of TIP60 complex residing within QTL-3d (*yeti* and *dRSF-1*) and three members outside QTL (*dom*, *E*(*Pc*) & *DMAP1*; **Supplemental table S6**) are differentially expressed between tolerant and sensitive genotypes [67,74,75].

Within QTL-21d, we did not find any genes with function in heterochromatin formation or DSB repair. However, the germline differentiation factor *brat* was exceptional in containing 14 in-phase SNPs in introns and downstream regions, and is upregulated in the tolerant genotypes (**Supplemental table S5** and **S11**). In adult ovaries, Brat is excluded from GSCs, but is expressed in CBs and promotes differentiation [76]. Because DNA damage blocks cystoblast differentiation by suppressing *bam* translation [21], *brat* could confer tolerance in older females by helping cytoblasts escape arrest.

### 6. Investigating the role of *brat* in tolerance

To determine the impact of *brat* on tolerance, we examined the tolerance phenotypes of a *brat* loss-of-function mutation *(brat^1^)* and multiple deficiencies overlapping *brat*. The candidate causative variants in *brat* that are proposed to influence tolerance are most likely heterozygous in dysgenic hybrid offspring. We therefore evaluated the heterozygous effect of *brat^1^* and overlapping deficiencies by comparing the incidence of ovarian atrophy between mutant or deficiency offspring to balancer siblings from dysgenic crosses *(brat/CyO* x Harwich).

In absence of dysgenesis, *brat* loss of function alleles impact oogenesis recessively [76]. However, we found that the *brat^1^* heterozygotes showed a significantly higher frequency of ovarian atrophy (68.6%) than their balancer control siblings (37.5%) (**Figure 6, Supplemental table S20**). Furthermore, two out of three deficiency stocks with deletions overlapping *brat* increased ovarian atrophy similarly to the *brat^1^* mutant, suggesting that this phenotype is not an effect of the 2^nd^ chromosome of the *brat^1^* mutant line (**Figure 6, Supplemental table S20**). The deficiency line (Df(2L)*brat* [ED1231]) that shows no change in the incidences of ovarian atrophy may carry deletions in genes with opposing function to that of *brat*, or suppressors elsewhere in the genome. Our results suggest that *brat* activity increases fertility in dysgenic females, which is consistent with our observation that tolerant alleles exhibit increased *brat* expression (**Supplemental table S5)**. Notably, the fertility effects of *brat* were observed in 3 day-old offspring, as attempts to look at older females (21 day-olds) were unsuccessful due to a high mortality rate.

**Figure 6.**
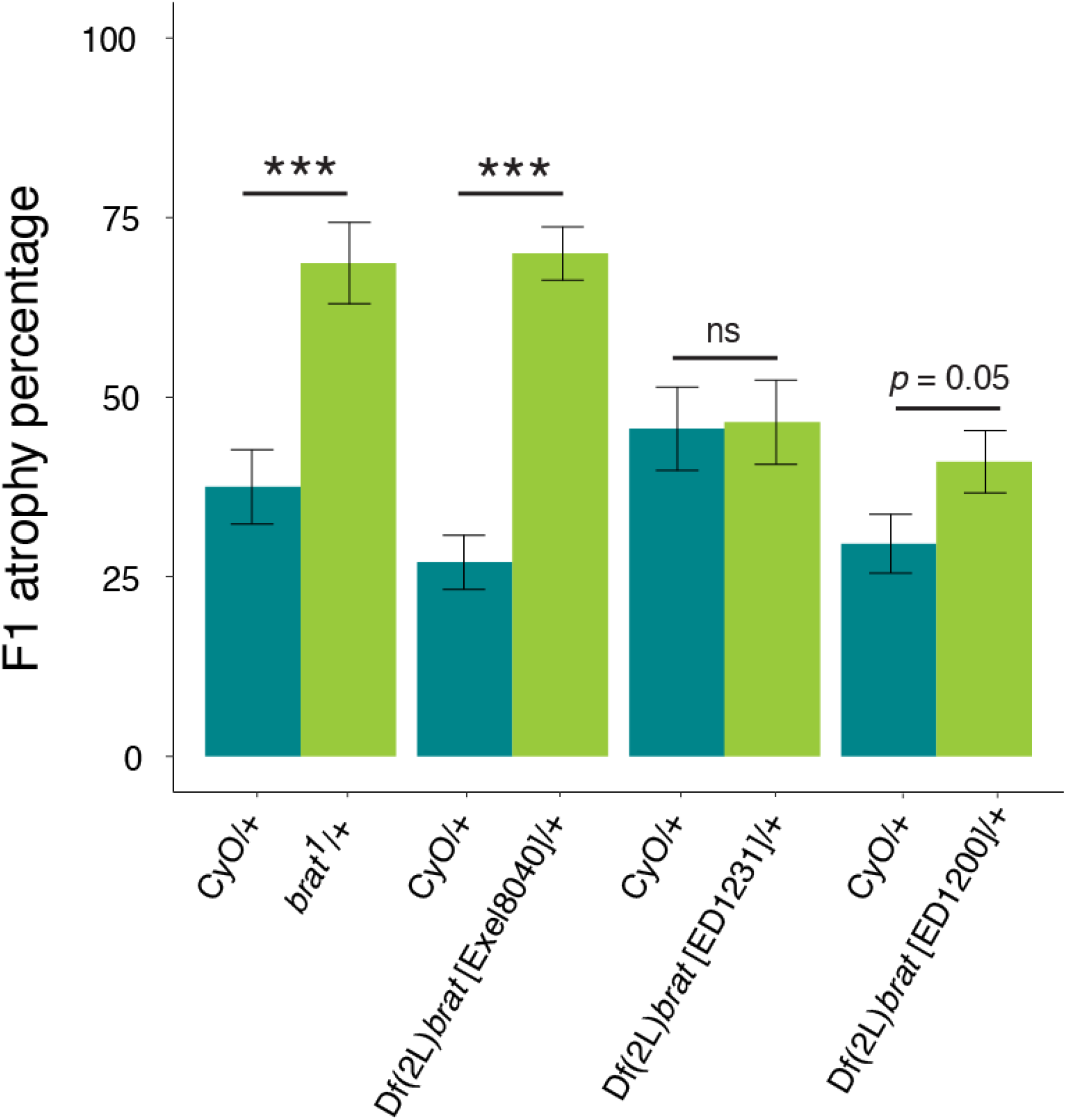
Loss-of-function mutation of *brat* increases severity of hybrid dysgenesis. The percentage of F1 ovarian atrophy is compared between control balancer siblings CyO/+, heterozygous *brat^1^* mutants and heterozygous deficiency lines Df(2L)*brat*. *brat^1^* mutant: χ^2^= 13.55, df=1, *p*-value =0.0002. Df(2L)*brat* [Exel8040]: χ^2^= 14.78, df=1, *p*-value =0.0001. Df(2L)*brat* [ED1231]: χ^2^= 0.06, df=1, *p*-value =0.8. Df(2L)*brat* [ED1200]: χ^2^= 3.66, df=1, *p*-value =0.05. The underlying data are provided in **Supplemental table S20**.

## Discussion

Although small RNA mediated TE regulation is widely studied, little is known about cellular and molecular mechanisms that confer tolerance to transposition. Here we uncovered natural variation in tolerance to *P*-element DNA transposons, which is associated with two or more loci proximal to the second chromosome centromere in *D. melanogaster*. We further showed that tolerant and sensitive genotypes may differ in their ability to enact DSB repair, potentially explaining their differential responses to *P*-element transposition. Finally, we identified candidate genes in each QTL that potentially determine the phenotypic differences between tolerant and sensitive alleles. Within QTL-3d, *Nipped-A* has a non-synonymous in-phase SNP that could alter the activity of encoded protein. By contrast, *brat*, located in QTL-21d, has in-phase SNPs in its intronic and downstream regions, and is upregulated in tolerant genotypes.

### Differences in DSB repair and TIP60 activity

We propose that in young females, tolerance is determined by the ability to repair DSBs resulting from *P*-element activity in larval PGCs. In our ovarian RNA-seq data, we saw two circumstantial indicators of differences in DSB repair in tolerant genotypes: increased chorion gene expression and decreased histone gene expression (**Figure 2b-d**). Chorion gene amplification is dependent upon DSB repair; thus, while we did not directly assay amplification, their increased expression may indicate more efficient repair [34]. Conversely, in yeast, excess histones inhibit DSB repair, potentially by competing with repair complexes for access to DNA [40,77]. Increased histone expression in sensitive genotypes may therefore inhibit DSB repair. Consistent with both of these observations, we observed that tolerant genotypes are significantly more resilient to X-ray radiation (**Figure 3**), which is widely associated with increased activity of DNA repair genes [51–55].

Enhanced repair in tolerant genotypes may be explained by increased activity of the TIP60 complex: a conserved chromatin remodeling complex with functions in DSB repair [75,78] and heterochromatin formation [66–69]. While two additional TIP60 components reside within QTL-3d and are differentially expressed between sensitive and tolerant genotypes (*yeti* and *dRSF-1), Nipped-A* is unique in containing a non-synonymous in-phase SNP. Consistent with a deleterious effect, the amino acid change carried by the sensitive allele is quite rare in recently sampled natural populations worldwide (collected after *P*-element invasion), occurring in only four of 645 sequenced strains [79,80]. Interestingly, one of these strains (RAL799) was recently examined for radiation sensitivity and found to be highly sensitive [81].

While the functional consequences of the non-synonymous SNP that separates tolerant and sensitive *Nipped-A* alleles is not clear, the upregulation of four other TIP60 members in tolerant genotypes (*dRSF-1, dom, E(Pc*) & *DMAP1*), together with evidence of enhanced heterochromatin formation, suggests increased TIP60 activity (**Supplemental table S6**). Increased TIP60 could directly facilitate DSB repair through its function in the exchange of phosphorylated Histone 2AV at DSBs [75]. However, enhanced heterochromatin formation resulting from TIP60 function could also facilitate DSB repair indirectly by reducing the expression of histones. The latter is more speculative, because although the histone locus body has a specialized chromatin state determined by multiple suppressors of variegation [43,45], there is limited evidence that TIP60 regulates the histone locus body [82].

### Germ cell differentiation and tolerance in adult females

We identified *brat* as a promising candidate to explain natural variation in tolerance of aging (21 day) females. *brat* resides in QTL-21d, contains 14 in-phase SNPs and was upregulated in tolerant genotypes. Consistent with *brat* function promoting tolerance, we observed that a *brat* loss-of-function mutation and multiple *brat* deficiencies are dominant enhancers of dysgenic ovarian atrophy (**Figure 6b**), while their effects on oogenesis in non-dysgenic germlines are recessive [83].

We propose that in aging adult females, *brat* could confer tolerance by promoting cystoblast (CB) differentiation, thereby opposing the arrested differentiation that results from DSBs [Figure 7b; 21]. Indeed, CB accumulation is observed when hybrid dysgenesis is induced by temperature shift in adult females [15,20]. CB differentiation is delayed following DNA damage by repressing the translation of *bam*, a key differentiation factor [21]. Brat acts to promote the translation of *bam* by repressing the translation of Mad and Myc [83]. Interestingly in the larval gonad, Myc activity is associated with retention of PGCs in dysgenic germlines [17], further highlighting how tolerance mechanisms may differ over the course of development.

**Figure 7.**
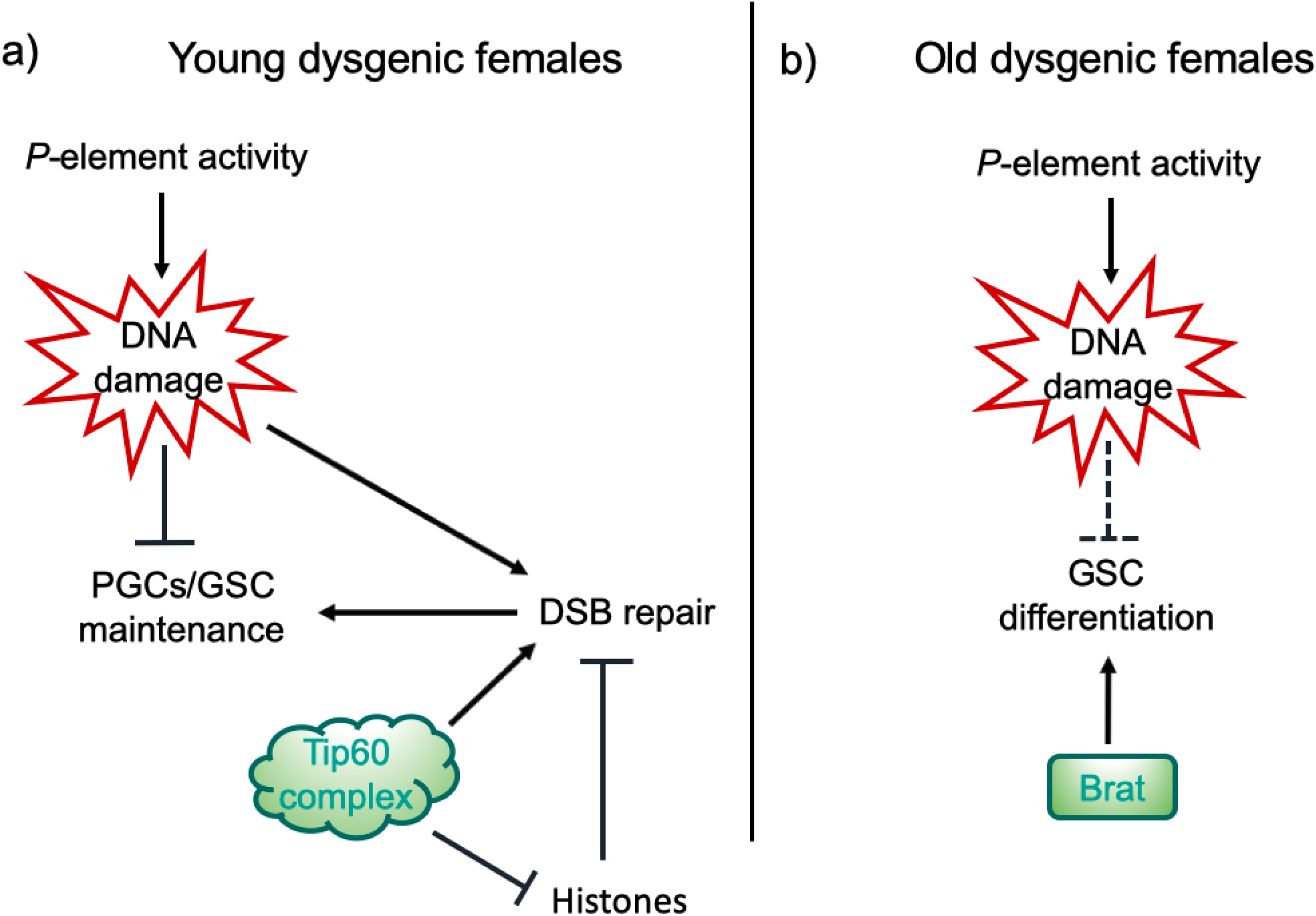
Hypothesized mechanisms of TE tolerance in young and old females. a) We propose that in larval and young-adult females, germline tolerance to *P*-elements may be determined by enhanced DSB repair through increased TIP60 activity. b) In aging dysgenic females, *brat* may determine tolerance by promoting differentiation of arrested cytoblasts, thus aiding in their escape from the cell-cycle arrest imposed by *P*-element mediated DNA damage.

Our demonstration that *brat* promotes tolerance also presents an intriguing contrast to our previous report that another germline differentiation factor, *bruno*, reduces tolerance [14]. Unlike *brat* alleles, *bruno* alleles and deficiencies are dominant suppressors of hybrid dysgenesis. Similar to *brat, bruno* encodes an mRNA binding protein that promotes differentiation of pre-meiotic cysts in the female germline, albeit at a later 4-cell stage [84,85]. *bruno* further differs from *brat* in acting independently of the Bam/Bgcn pathway that is repressed in CBs after DSBs [21,86]. Collectively therefore our data speak to a careful orchestration of germ cell differentiation that can facilitate germline persistence in the face of DNA damage.

### Conclusion

Our work reveals that natural tolerance to transposition can arise throughout the lifecycle of the fly, ensuring the maintenance of germline cells during development and the production of gametes in adults. This contrasts our previous study of natural variation in the population A RILs of the DSPR, which uncovered a single major effect QTL of differentiation factor, *bruno*, on tolerance in both young and old females [14]. Furthermore, while DNA damage signaling is a clear determinant of dysgenic germ cell loss [15,19,20], we for the first time provide evidence of natural variation in DNA repair offsetting the damaging effects of transposition. Our observations therefore point to multiple new mechanisms through which germlines could withstand the genotoxic effects of unregulated transposition, which may respond to natural selection after new TEs invade.

## Methods

### *Drosophila* Strains and Husbandry

The recombinant inbred lines (RILs) were generously provided by Stuart Macdonald. Harwich (#4264), *b cn* (#44229), *brat^1^* (#3988), Df(2L)*brat*[Exel8040] (#7847), Df(2L)*brat* [ED1231] (#9174) and Df(2L)*brat* [ED1200] (#9173) were obtained from the Bloomington *Drosophila* stock center. Canton-S was obtained from Brigitte Dauwalder. All flies were maintained in standard cornmeal media.

Alleles of the second chromosome centromeric region, containing both QTL, were extracted from three recombinant inbred lines carrying B6 QTL allele (#21076, #21218, #21156) and two RILs carrying B8 QTL allele (#21077, #21154) into a common background by crossing them to multiply marked stocks *b cn* (#44229). After 7 rounds of backcrossing followed by inbreeding, the final isogenic lines (Sensitive_B6^1^, Sensitive_B6^2^, Sensitive_B6^3^ and Tolerant_B8^1^, Tolerant_B8^2^) were generated. The lines were made homozygous for the 2^nd^ chromosome by inbreeding and selecting for wild type phenotype. The genotype of the isogenic lines were verified through PCR using five different primers within the two QTL. chr2L:19383155-19383970: AACCCTTTTTCGCTGACAATAACA, ATTATCAGCAGGAGCCGGAAACTT; chr2L:21333500-21334300: AAGTGAAGCTAACAACGTGACAAC,CGTTTGACCATCGCTTACAACTAA; chr2R:2392800-2393600: AACAGGAGGTCGAAAGCCAAATA, ATGCAGAGTCATATTCTGGGTTGG; chr2R:6203290-6204284: AATGGAGACCGTTGATTTTGGTAA,CTTTTCTGCGGCATCAGGTG; chr2R:6058000-6059000: TGGCAATTGCAATCCTTTTGGTAT, ATAACACGAACTACGACCTTTCCA

### Phenotyping

Phenotyping of ovarian atrophy was performed as described previously in Kelleher *et al* [14]. Briefly, crosses between virgin RIL females and Harwich males were transferred to fresh food every 3-5 days. Since crosses reared at a restrictive temperature (29 °C) result in complete gonadal atrophy in F1 offspring, we reared our crosses at a lower permissive temperature (25 °C), which produces an intermediate phenotype that better reveals the variation in severity of dysgenesis [12,14,15,87]. F1 offspring were maintained for 3 days or 21 days, at which point their ovaries were examined using a squash prep [87]. 21 day-old females were transferred onto new food every 5 days as they aged to avoid bacterial growth. Females who produced 1 or more chorionated egg chambers were scored as having non-atrophied ovaries, and females producing 0 egg chambers were scored as having atrophied ovaries.

Crosses and phenotyping were performed for 673 RILs across 22 experimental blocks for 3 day-old F1 females, and 552 RILs across 18 experimental blocks for 21 day-old F1 females. If fewer than 21 F1 offspring were phenotyped for the same cross, it was discarded and repeated if possible. In total, we phenotyped >20 3-day old and 21 day-old F1 female offspring for 595 RILs and 456 RILs, respectively.

### QTL mapping

QTL mapping was performed as described in Kelleher *et al*. [14]. Briefly, for each developmental time point, we modeled the arcsine transformed proportion of F1 ovarian atrophy as a function of two random effects: experimental block and undergraduate experimenter. Regression models were fit using the lmer function from the lme4 package [88]. We then used the residuals as a response for QTL mapping with the DSPRqtl package [22] in R 3.02 [89]. The LOD significance threshold was determined from 1,000 permutations of the observed data, and the confidence interval around each LOD peak was identified by a difference of −2 from the LOD peak position (Δ2-LOD) [25], or from the Bayes Confidence Interval [90]. For Δ2-LOD intervals, we took the conservative approach of determining the longest contiguous interval where the LOD score was within 2 of the peak value. We further calculated the broad sense heritability of ovarian atrophy as in Kelleher *et al*. [14].

### Estimation of Founder Phenotypes and QTL phasing

To estimate the phenotypic effect associated with each founder allele at the QTL peak, we considered the distribution of phenotypes from all RILs carrying the founder haplotype at the LOD peak position (genotype probability >0.95%) [22]. QTL were then phased into allelic classes by identifying the minimal number of partitions of founder haplotypes that describes phenotypic variation associated with the QTL peak, as described previously [14,22].

### Fertility Assays

Virgin female offspring from dysgenic crosses between isogenic lines carrying tolerant_B8_1_/B8_2_(21077, 21154) and tolerant_B8_2_/B8_3_ (21218, 21156) alleles and Harwich males were collected daily and individually placed in a vial containing two Canton-S males. Females were allowed to mate for 5 days and were transferred to a new vial for another 5 days after which the parents were discarded. The presence and total number of F2 individuals were counted from the two vials.

### Identification of in-phase polymorphisms

The SNP data of B founders that used to infer in-phase SNPs is based on dm3 [22]. To identify in-phase SNPs we looked for alternate SNP alleles that match the predicted phenotypic class for each of the QTL peaks. For QTL-21d we used the criteria: sensitive class (B2, B6) and the tolerant class (B1, B3, B4, B7, B8), whereas for QTL-3d: sensitive class (B6) and the tolerant class (B1, B2, B3, B4, B7, B8).

### Selection of paired RILs with alternate QTL alleles

We identified background matched RILs containing either the B6 (“sensitive”) or B4 (“tolerant”) haplotypes from the start position of the QTL-21d confidence interval (2L: 19,010,000) to the end position of QTL-3d confidence interval (2R: 6,942,495) (*P* > 0.9), based on their published HMM genotypes [22]. For all possible RIL pairs (B6 and B4), we then calculated the number of 10 Kb genomic windows in which they carried the same RIL haplotype (*P* > 0.9). We selected three pairs of RILs, which carry the same founder genotype for 47% (21213 & 21183), 46% (21147 & 21346) and 44% (21291 & 21188) of genomic windows outside of the QTL.

### Small RNA-seq and total RNA-seq

RILs were maintained at 25°C, and three biological replicates of 20 ovaries were dissected from 3-5 day old females. Ovaries were homogenized in TRIzol and stored at −80°C until RNA extraction. 50 μg of total RNA from each of 18 biological samples (3 biological replicates x 3 pairs) was size fractionated in a 15% denaturing polyacrylamide gel and the 18-30 nt band was excised. 2S-depleted small RNA libraries for Illumina sequencing were then constructed according to the method of Wickersheim and Blumenstiel [91]. Ovarian small RNA libraries were published previously [SRP160954, 92]. Ribodepleted and stranded total RNA libraries were generated from the same ovarian samples using NuGen total RNA kit (TECAN). All 18 small RNA and total RNA libraries were sequenced on an Illumina Nextseq 500 at the University of Houston Seq-N-Edit Core, and are deposited in the NCBI BioProject PRJNA490147.

### Small-RNA analysis

Sequenced small RNAs were separated based on size into miRNAs/siRNAs (18-22nt) and piRNAs (23-30nt) [11]. Reads corresponding to contaminating rRNAs, including 2S-rRNA, were removed from each library by aligning to annotated transcripts from flybase [93]. To determine the piRNA cluster activity we first uniquely aligned the piRNAs to reference genome (dm6 [28]) using Bowtie1 (-v 1 -m 1) [94]. We then used a customized perl script (https://github.com/JLama75/piRNA-cluster-Coverage-script) to count reads that mapped to a set of previously annotated piRNA clusters from the same genotypes (497 piRNA clusters, [95]). Read counts normalized to total mapped microRNAs for each library were used to infer differential expression using DESeq2 [96]. Sliding window estimates of piRNA abundance (**Figure 2c and d)** were calculated using bedtools genomecov [97], normalizing the read counts to total mapped miRNA reads.

### Total RNA analysis

Residual ribosomal RNAs (rRNAs) were identified in ribodepleted libraries based on alignment to annotated rRNAs from flybase [93], and excluded from further analysis. Retained reads aligned to the library of consensus satellite and TE sequences from repbase [98], plus additional satellite consensus sequences from Larracuente [99]. For TE expression, the total reads mapped to TE sequences were counted using unix commands (uniq -c). Remaining reads that failed to map were aligned to *D. melanogaster* transcriptome (dm6/BDGP6) using Kallisto with default parameters [100]. Differentially expressed TEs and genes were identified from a combined analysis in DESeq2 [96]. Genes and TEs with base mean >= 100, Adjusted *P*-value <= 0.05 and whose expression pattern differed (fold change >= 1.5) were considered differentially expressed between the B6 and B4 QTL haplotype.

### Radiation Sensitivity

Third instar larvae were either mock treated or irradiated in a Rad Source RS 1800 X-ray machine set at 12.5 mA and 160 kV. To obtain 3rd instar larvae, embryos were collected for 24 hr and aged for 5 days at 25 degree Celsius. The food vials containing larvae were then X-ray irradiated at doses from 5-80 Gray after which an optimal dose that clearly depicts the phenotypic difference was selected. Survival to adulthood was determined by scoring the number of empty and full pupal cases at 10 days after radiation.

---

**Figure S1).**
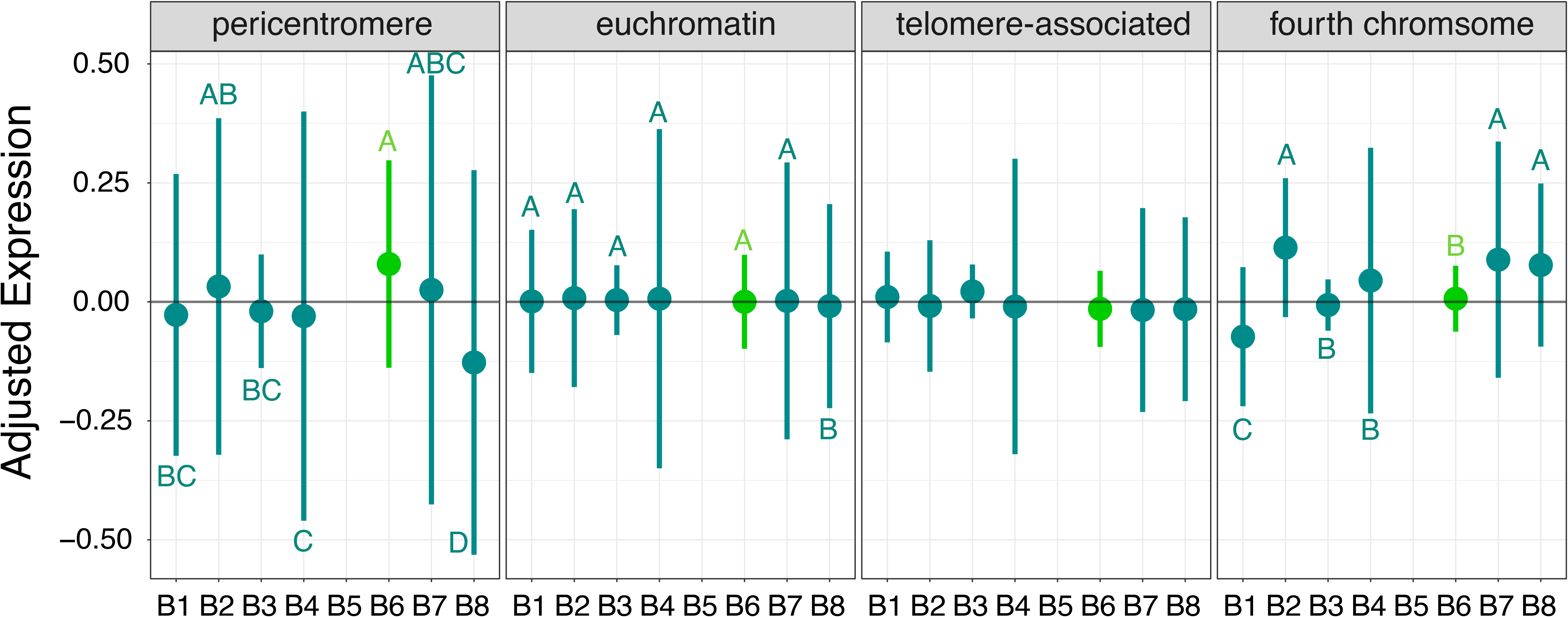
Sensitivity is associated with increased expression of pericentromeric genes in the head. a) Mean expression of genes located in the pericentromere, euchromatin, telomere and the fourth chromosome from RILs carrying each of the eight B founder genotypes at the QTL-3d region. Error bars represent the standard deviation among mean expression levels of different genes. The sensitive/B6 (light green) shows high pericentromeric gene expression compared to the tolerant strains (dark green) (Anova; *F_6,494_*=7.775, P<5.24e-08). The letters indicate significantly different expression levels based on Tukey-HSD comparisons between RILs with different founder alleles.

**Figure S2).**
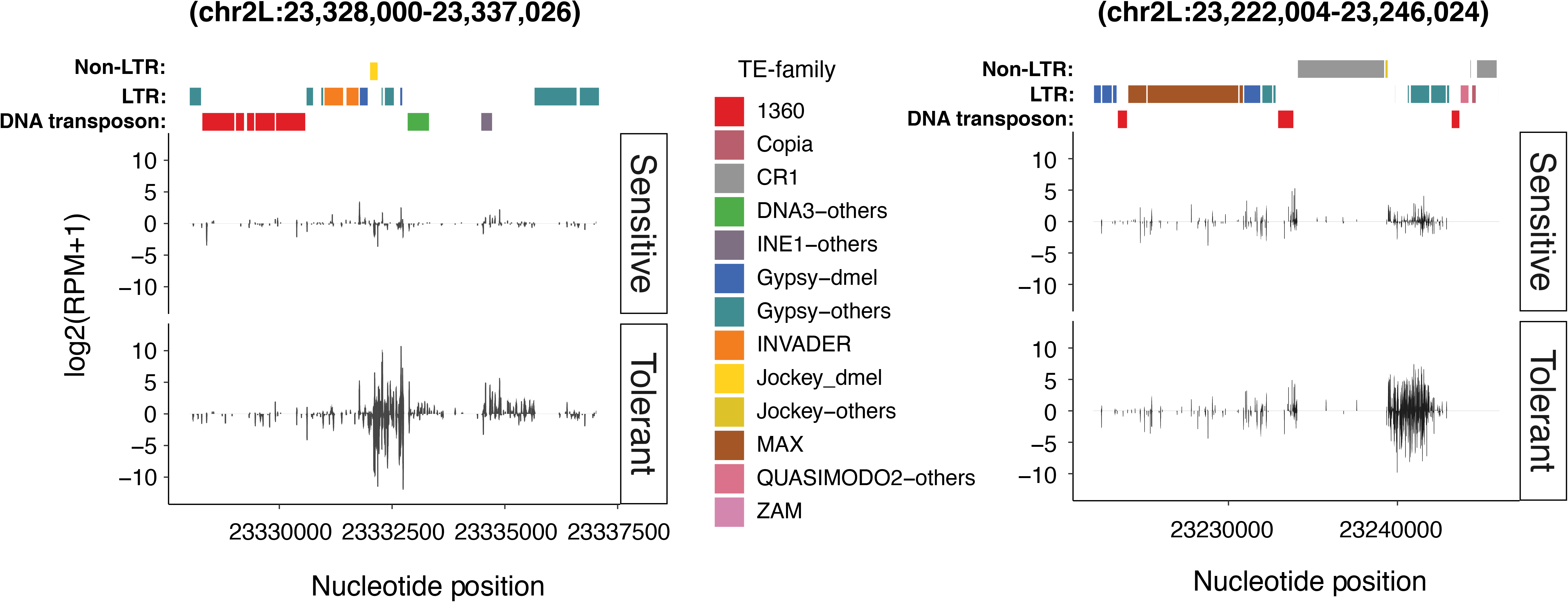
Expression profile of QTL piRNA clusters in a sensitive and tolerant NIL pair. The piRNA expression between sensitive and tolerant genotypes from 21188-21291 NIL pairs along the two QTL piRNA clusters: 2L:23,328,000-23,337,026 and 2L:23,222,004-23,246,024, respectively. Only uniquely mapping piRNAs are considered. The TE families at the top of each panel are represented by different colors. TE-others represent the repeat families coming from sibling species of D. melanogaster. Positive value indicates piRNAs mapped to the sense strand of the reference genome and negative value indicates those from the antisense strand. The piRNA cluster expression levels are estimated by log2 scale transformed of reads per million mapped reads [log2(RPM+1)].

**Figure S3).**
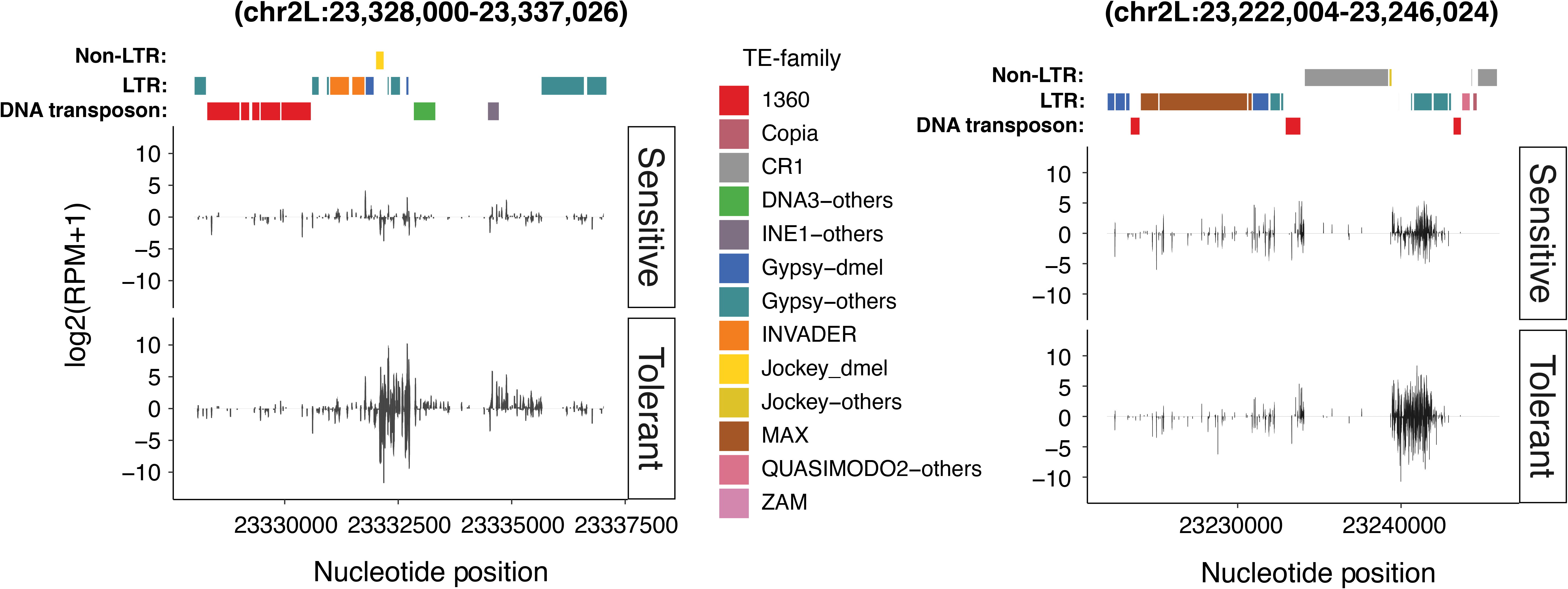
Expression profile of QTL piRNA clusters in a sensitive and tolerant NIL pair. The piRNA expression between sensitive and tolerant genotypes from 21346-21147 NIL pairs along the two QTL piRNA clusters: 2L:23,328,000-23,337,026 and 2L:23,222,004-23,246,024, respectively. Only uniquely mapping piRNAs are considered. The TE families at the top of each figure are represented by different colors. TE-others represent the repeat families coming from sibling species of D. melanogaster. Positive value indicates piRNAs mapped to the sense strand of the reference genome and negative value indicates those from the antisense strand. The piRNA cluster expression levels are estimated by log2 scale transformed of reads per million mapped reads [log2(RPM+1)].

**Figure S4.**
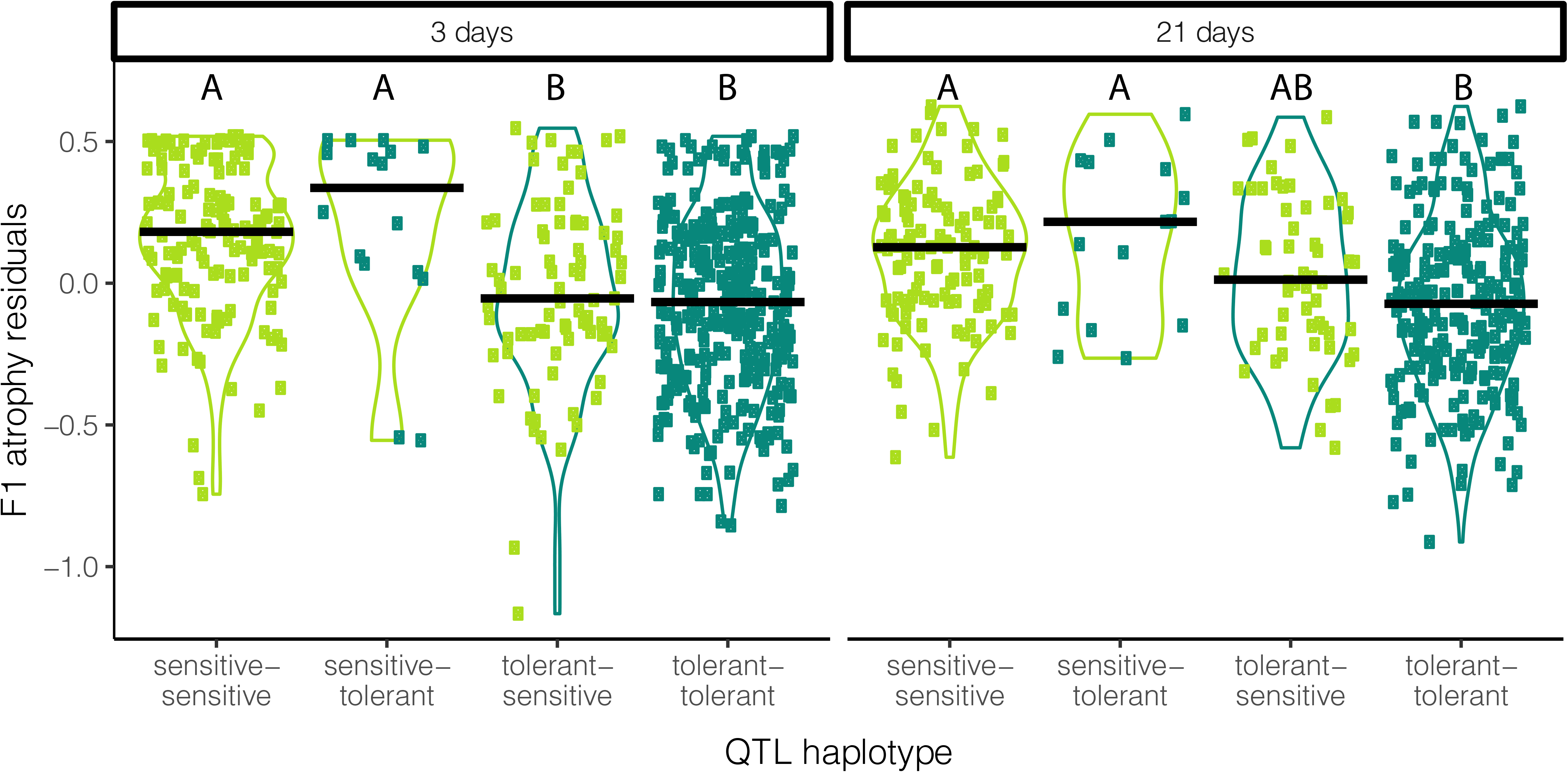
Tolerance among 3 and 21 day females based on QTL haplotype. Four haplotypes are compared, which comprise all possible combinations of tolerance alleles at 2 QTL. The allele at the 3 day QTL is indicated first and is represented by the color of the violin plot (light green = sensitive, dark green = tolerant). The allele at the 21 day QTL is indicated second and represented by the color of the points on the scatter plot. Y-axis is residual variation in F1 atrophy after accounting for student experimenter and block. Among 3 day old females, haplotypes containing different alleles for the 3 day old QTL are significantly different from each other (Tukey HSD P=0.016-0). However, haplotypes containing alternative QTL for the 21d only do not differ from each other (Tukey HSD P>0.74). This suggests phenotypic variation in 3 day old females is not influenced by their genotype at the 21 day QTL. In contrast, among 21 day old females tolerant alleles in both QTL loci are required to significantly increase tolerance above sensitive allele containing haplotypes (Tukey HSD *P* = 0.01-0).

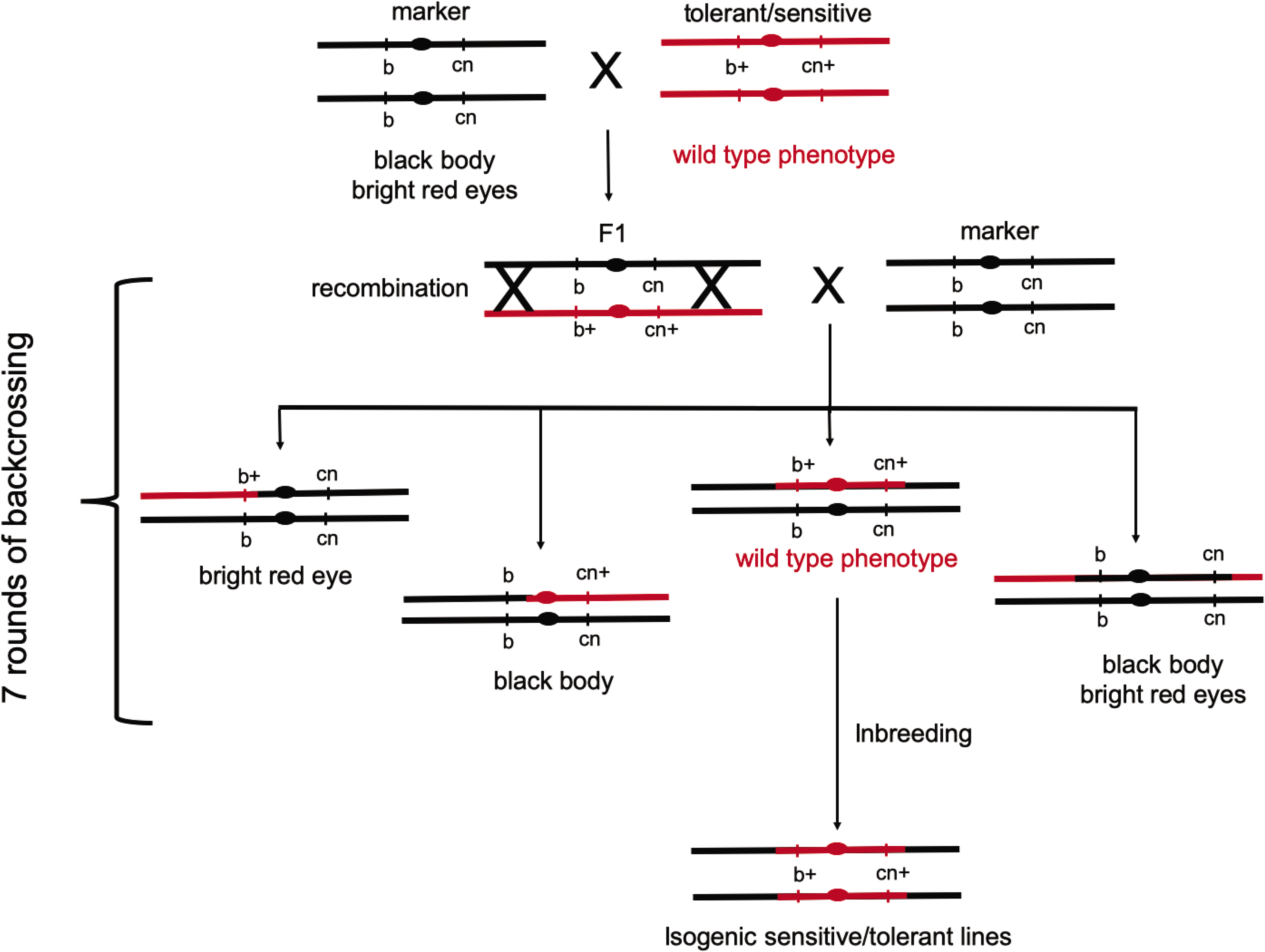

